# Monte Carlo simulations predict distinct real EEG patterns in individuals with high and low IQs

**DOI:** 10.1101/2025.01.07.631738

**Authors:** Arturo Tozzi

## Abstract

The neural mechanisms underlying individual differences in intelligence are a central focus in neuroscience. We investigated the effectiveness of Monte Carlo simulations in predicting real EEG patterns and uncovering potential neural differences between individuals with high and low intelligence. EEG data were collected from two groups of volunteers categorized by IQ, namely, a high-IQ group and a low-IQ group. A univariate normal distribution was fitted to each EEG channel using Maximum Likelihood Estimation, after which synthetic datasets were generated based on the estimated parameters. Statistical analyses including Root Mean Square Error (RMSE) calculations assessed the alignment between real and simulated data. We showed that Monte Carlo simulations effectively replicated the statistical properties of the EEG data from both the groups, closely matching the real central tendencies, variability and overall distribution shapes. Specific EEG channels, particularly in the frontal and temporal bilateral regions, exhibited significant differences between the two groups, pointing to potential neural markers of cognitive abilities. Further, the low-IQ group exhibited higher predictability and more consistent neural patterns, reflected by lower RMSE values and smaller standard deviations across several EEG channels. Conversely, the high-IQ group displayed greater variability and larger RMSE values, reflecting complex neural dynamics that are less predictable by Monte Carlo simulations. Our findings underscore the utility of Monte Carlo simulations as a robust tool for replicating EEG patterns, identifying cognitive differences and predicting EEG activity associated with intelligence levels. These insights can inform predictive modeling, neurocognitive research, educational strategies and clinical interventions of targeted cognitive enhancement.

## INTRODUCTION

Exploring the neural mechanisms underpinning intelligence has been a longstanding primary focus of cognitive neuroscience research. Electroencephalography (EEG) offers unique insights to assess differences in cognitive abilities, including distinctions between individuals of varying intelligence levels (Friedman et al., 2019). With its high temporal resolution, the non-invasive EEG evaluates the interplay between synchronization, complexity and network efficiency (van Dellen et al., 2015). For instance, higher IQ is associated with reduced long-distance EEG information flow and enhanced local processing efficiency, supporting small-world models (Thatcher et al., 2016). Short EEG phase delays and increased coherence in frontal regions correlate with higher intelligence, emphasizing the role of frontal lobe synchronization (Thatcher et al., 2005). Resting-state EEG studies have further explored intelligence-related differences, reporting balanced inter-hemispheric coordination in alpha and beta bands in more intelligent individuals (Jahidin et al., 2013). Also, it has been demonstrated that IQ correlates negatively with EEG energy but positively with information flow intensity at specific frequencies, emphasizing the role of efficiency in neural communication (Luo et al., 2021). Changes in microstate dynamics are associated with fluid intelligence and its enhancement following cognitive training (Santarnecchi et al., 2017). Lu et al. (2022) found that individuals with higher fluid intelligence allocate attentional resources more flexibly, particularly in complex tasks, as reflected in theta and alpha EEG activities. Together, these findings underscore the utility of EEG in the assessment of the neuronal mechanisms of intelligence, revealing consistent patterns of neural efficiency, inter-hemispheric coordination and adaptive resource allocation.

Conversely, the analysis of EEG data poses significant challenges due to their inherent variability, high dimensionality and sensitivity to noise (Hassani et al., 2015). To address these challenges and enhance our ability to model and predict EEG patterns, advanced statistical and computational methods are required. Monte Carlo simulations have been widely used across various scientific disciplines, providing a powerful framework for modeling complex systems influenced by variability and uncertainty (Metropolis and Ulam, 1949; Rubinstein and Kroese, 2016). By leveraging statistical properties derived from observed data, Monte Carlo simulations generate synthetic datasets that may reflect real-world behaviors (Salvadori et al., 2024; Jones and Fleming, 2024). A Monte Carlo approach could be particularly well-suited for EEG data, as it allows researchers to explore and replicate neural dynamics without the need for extensive experimental data collection. Monte Carlo methods have been applied in neuroscience to simulate and analyse electromagnetic brain signals, providing approximations of event-related brain activity (Herdman 2021). Monte Carlo simulations have been utilized for brain source localization (Georgieva et al., 2013) and evaluation of errors as a function of position within the brain in MRI-MEG/EEG co-registration techniques (Singh et al., 1997). Surprisingly, it has been demonstrated that EEG localization is more accurate than MEG localization for the same number of sensors averaged over many source locations (Liu et al., 2002). Monte Carlo analysis can also simulate event-related changes in amplitude and phase-amplitude correlations, enabling close approximations of real EEG and MEG data (Herdman 2021). This approach is particularly valuable for validating data analysis methods, including measurements of functional connectivity and phase-amplitude coupling. A Bayesian framework has been introduced for parameter estimation in EEG modeling using a marginalized Markov Chain Monte Carlo approach (Hettiarachchi et al., 2012). This method was employed to fit a neural mass model to EEG data effectively.

Despite these studies, the application of Monte Carlo simulations to EEG research in revealing cognitive differences between individuals of high and low intelligence remains relatively underexplored. This study aims to bridge this gap by investigating how Monte Carlo simulations can model, simulate and reproduce real EEG traces. Central to this investigation is the issue of predictability: using Monte Carlo simulations, are the EEG patterns from high-IQ individuals more or less predictable than those of their lower-IQ counterparts? It can be hypothesized that higher cognitive abilities are associated with greater neural flexibility and variability, potentially reducing predictability in simulations. Conversely, lower intelligence may correspond to more stable neural patterns, increasing predictability. Monte Carlo simulations, by generating synthetic EEG data modeled on the statistical properties of real datasets, offer a systematic approach to testing these hypotheses.

We conclude that that Monte Carlo methods are a robust tool for exploring the neural dynamics of intelligence, paving the way for future investigations into brain-behavior relationships. By accurately replicating EEG patterns and identifying significant group differences, Monte Carlo simulations contribute to a deeper understanding of the cognitive and neural processes underlying intelligence.

## MATERIALS AND METHODS

### Participants and Data Collection

This study retrospectively builds on the foundational research conducted by Norbert and Ksenija Jaušovec through 2010 (Jaušovec and Jaušovec, 2001; 2003; 2005; 2010), which was later advanced in collaboration with Tozzi et al. (2021a; 2021b). This continuation of their work is undertaken with great respect and recognition of Norbert’s untimely passing.

EEG data were collected from two groups of right-handed volunteers categorized by IQ, namely, a high-IQ group and a low-IQ group. Each group consisted of five participants, yielding a total sample of 10 individuals (mean age: 19.8 years; SD = 0.9; range = 18–21 years; males: 4). The IQ categorization was based on standardized test scores, with the high-IQ group representing the top quartile (IQ SD = 127) and the low-IQ group representing the bottom quartile (IQ SD = 87). Differences in EEG activity between these groups were analyzed during the performance of two oddball tasks (auditory and visual). The study adhered to the Declaration of Helsinki and received approval from the Ethics Committee of the University of Maribor, Slovenia.

EEG signals were recorded using a 64-channel system to ensure comprehensive cortical coverage. Electrode placement followed the 10–20 international system, covering nineteen scalp locations: [FP1], [FP2], [F3], [F4], [C3], [C4], [P3], [P4], [O1], [O2], [F7], [F8], [T3], [T4], [T5], [T6], [CZ], [FZ] and [PZ]. The electrodes were sintered Silver/Silver Chloride (8mm diameter) with impedance maintained below 5 kΩ. All leads were referenced to linked mastoids (A1 and A2), with a ground electrode on the forehead. Vertical eye movements were recorded using additional electrodes placed above and below the left eye. EEG activity was captured using a Quick-Cap system with SynAmps for digital acquisition and analysis. Signals were digitized at 1000 Hz with a gain of 1000 (resolution: 0.084 μV/bit, 16-bit A/D conversion) and stored on a hard disk. Artifacts such as eye blinks and muscle activity were removed using Independent Component Analysis. Data were then band-pass filtered between 1 and 40 Hz to isolate relevant neural activity while minimizing noise. The time-series data were located in separate columns, each one corresponding to a different EEG electrode. The data were numerical and represented EEG signal amplitudes in microvolts over time. Each row corresponded in the recording to a single time point in milliseconds.

Next, Monte Carlo simulations were performed to generate synthetic EEG traces, modeling the variations of future data points based on statistical distributions.

### Monte Carlo simulations

A reliable methodological framework was essential to ensure robust analysis and enabling meaningful comparisons between EEG traces of high-IQ and low-IQ individuals. The analysis of the EEG data followed a series of structured steps. The first step involved loading EEG datasets for both the groups. Multiple files for each group were then concatenated into unified datasets. Any rows or cells with missing values were removed to prevent inaccuracies in statistical calculations. The next step was distribution fitting, where statistical parameters, i.e., the mean μ and standard deviation σ, were derived for each EEG channel to simulate data closely reflecting real-world observations. Each channel was analyzed separately. A univariate normal distribution was fitted to the data using Maximum Likelihood Estimation (MLE) to calculate μ and σ, ensuring that the simulated data aligned with the distributions observed in the real datasets. The alignment of the fitted distributions were then validated by comparing histograms of real data against the simulated distributions.

Once the distributions were established, synthetic EEG traces were generated using Monte Carlo simulations. For each EEG channel, 100 random samples were drawn from the fitted normal distribution parameters μ and σ. The synthetic samples were organized into a structured dataset mirroring the original format, with separate columns for each channel. The simulation was performed independently for both high-IQ and low-IQ groups to preserve their distinct statistical properties. Following data simulation, statistical analyses evaluated the differences between real and simulated datasets and compared the predictability of the high-IQ and low-IQ groups. Fit accuracy was assessed by comparing real and simulated data using Root Mean Square Error (RMSE) for each EEG channel, where lower RMSE values indicated better alignment. Small adjustments were necessary to address mismatches in the number of samples between the real and the simulated data and to ensure data alignment during the RMSE calculation. Variability was also analyzed, as higher variability (e.g., larger standard deviations) in a dataset could reduce predictability by introducing greater spread around the mean. A larger number of EEG channels showing significant differences between real and simulated data suggested lower predictability. Two-sample t-tests and Welch’s t-tests were performed to identify significant differences between the real high-IQ and low-IQ groups. Visual comparisons, including histograms and boxplots, were produced to illustrate the alignment between real and simulated datasets.

All analyses and visualizations were performed using Python. Libraries such as numpy, pandas, and scipy were used for statistical calculations, while matplotlib was employed for creating visualizations. The scipy.stats module provided functions for t-tests and normal distribution fitting.

## RESULTS

Monte Carlo simulations effectively captured the central tendency, the spread and the variability of the original EEG traces. The real data distributions were closely approximated by the fitted normal distributions, indicating that the simulations accurately reflected the underlying statistical properties. The simulated variances closely aligned with those of the real data, staying within acceptable margins. Visual comparisons further highlighted the alignment between real and simulated data. Histograms demonstrated that the simulated data mirrored the shape and density of the real data, with only minor deviations observed in the tails for a few columns. Boxplots showed significant overlap in medians and interquartile ranges between the real and simulated data, underscoring the reliability of the modeling process. Moreover, the synthetic data successfully replicated the presence of outliers seen in the real dataset, further demonstrating its ability to reflect the inherent variability and complexity of the original data.

The Monte Carlo analysis revealed significant differences between the high-IQ and low-IQ groups in certain EEG channels (see **Table**). The t-tests and the Welch’s t-tests identified specific EEG channels where the groups exhibited statistically significant differences (p<0.05). Strong differences in activity levels were observed in left [FP1] and right [FP2] prefrontal regions. Channels [F3] and [F4] also showed significant differences in regions associated with higher-order cognition and decision-making. Central channels such as [CZ] exhibited marked differences, reflecting motor or cognitive integration, with [C3] and [C4] showing additional activity differences in motor-related regions. Temporal channels [T3], [T4] and [T5] demonstrated significant differences in regions linked to memory and language processing, while the occipital channel [O2] revealed distinctions in visual processing areas.

On the other hand, some EEG channels exhibited overlapping distributions between the groups, suggesting no significant differences in activity. These included [P3] and [PZ] in the posterior parietal regions, [T6] in the temporal region and [F7] in the left frontal area, where activity patterns appeared similar across groups.

In sum, significant differences between the high-IQ and low-IQ groups were found in medians and interquartile ranges for various channels. These findings suggest key areas where these groups differ in their distributions.

**Table.** Statistical Differences Across EEG Channels Between High-IQ and Low-IQ Groups.

| EEG Channel | p-Value | Significance |
| --- | --- | --- |
| [FP1] | 1.13e-20 | Significant |
| [FP2] | 3.11e-13 | Significant |
| [F3] | 1.13e-07 | Significant |
| [F4] | 8.86e-12 | Significant |
| [C3] | 8.28e-06 | Significant |
| [C4] | 3.77e-05 | Significant |
| [P3] | 0.741 | Not Significant |
| [P4] | 0.019 | Significant |
| [O1] | 0.558 | Not Significant |
| [O2] | 1.32e-05 | Significant |
| [F7] | 0.209 | Not Significant |
| [F8] | 4.78e-06 | Significant |
| [T3] | 0.002 | Significant |
| [T4] | 1.24e-19 | Significant |
| [T5] | 7.28e-16 | Significant |
| [T6] | 0.805 | Not Significant |
| [CZ] | 0.000368 | Significant |
| [FZ] | 1.33e-23 | Significant |
| [PZ] | 0.904 | Not Significant |

Visual comparisons provided further insights. The histograms highlighted distinct peaks or shifts between the two groups in significant channels, aligning with the statistical tests and confirming the observed differences (**Figure A**). The boxplots showed that the high-IQ group data had larger variability in significant channels, reflected in wider interquartile ranges, while the low-IQ group displayed more consistent and narrower distributions (**Figure B**). These patterns suggest that significant channels may serve as neural markers differentiating cognitive abilities between the groups.

The comparison of the predictability between the high-IQ and low-IQ groups revealed distinct insights (**Figure C**). RMSE values were generally lower for the low-IQ group, indicating better alignment between real and simulated data and suggesting higher predictability in the EEG traces of less intelligent individuals. Standard deviations were also slightly smaller for the low-IQ group across several EEG channels, further supporting its greater predictability. In contrast, the high-IQ group exhibited greater variability and larger RMSE values, which are indicative of patterns of reduced predictability in Monte Carlo simulations.

**Figures A-B.**
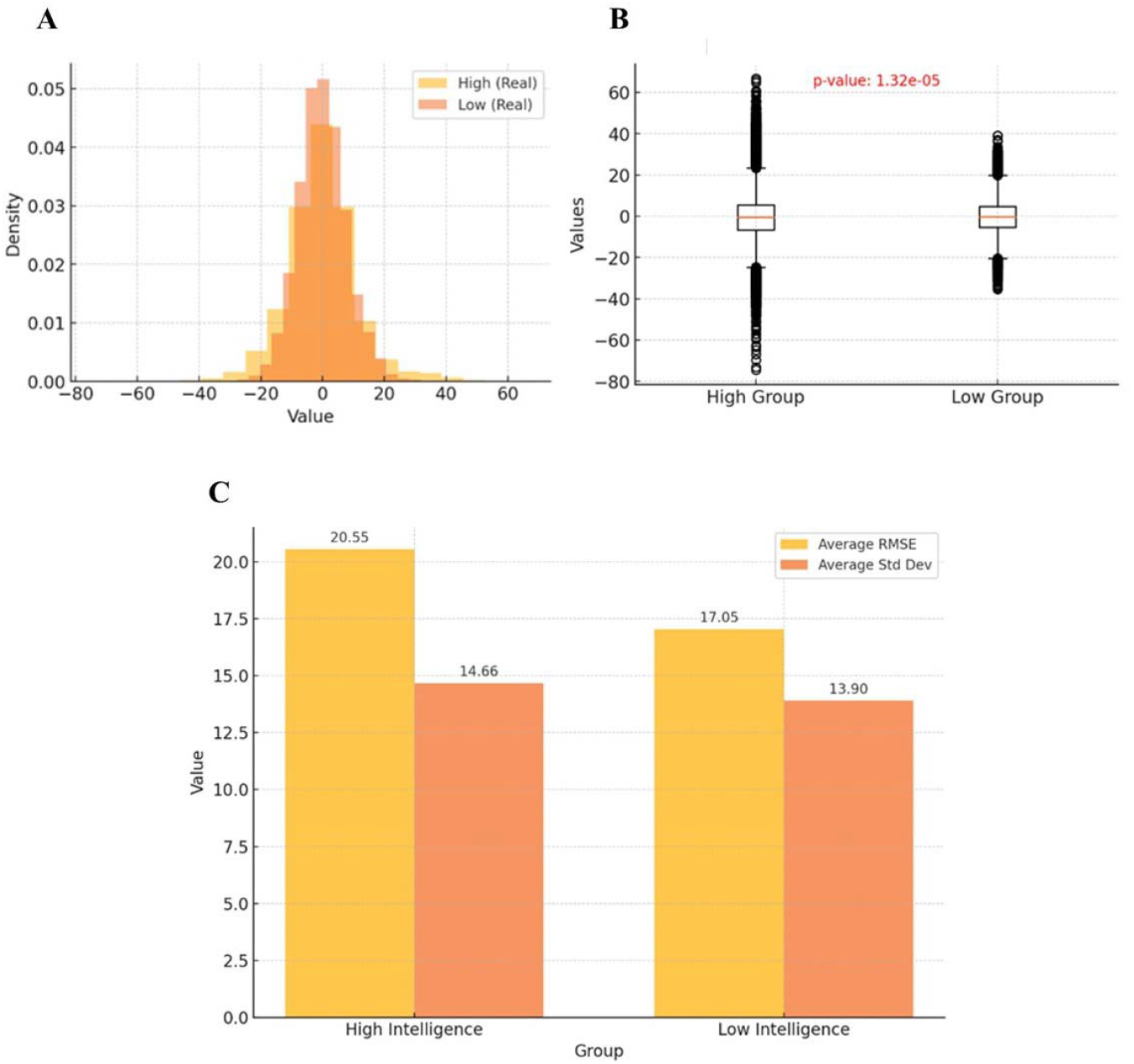
A visual comparison of Monte Carlo simulations showcasing the graphical representation of a single electrode [O2] as a representative example from the set of 19 electrodes. The histogram comparison (**Figure A**) highlights the differences in real distribution between high-IQ and low-IQ groups across each column. The corresponding boxplot (**Figure B**) visually compares the distributions of the two groups across each column. Differences in medians, interquartile ranges and potential outliers can be identified. Statistical significance is determined based on p-values obtained through t-tests. **Figure C**. Predictability of EEG traces via Monte Carlo simulation. This panel indicates that the EEG behaviour of the low-IQ group is more predictable, as reflected by their lower average RMSE and slightly smaller standard deviation compared to the high-IQ group.

## DISCUSSION

The results of this study underscore the remarkable utility of Monte Carlo simulations in modeling EEG traces and identifying significant differences between high-IQ and low-IQ groups. By accurately replicating the statistical properties of the original dataset, the simulated data closely mirrored the observed real patterns in terms of central tendencies, variability and overall distribution shapes. Our analysis revealed pronounced statistical distinctions in specific EEG channels, which suggest potential markers of cognitive ability. For instance, the differences observed in frontal regions [FP1] and [FP2] align with their roles in executive functions, attention and problem-solving. The high-IQ group exhibited greater variability in these areas, possibly reflecting more dynamic or complex neural processes. The temporal regions [T3] and [T4] showed disparities that could indicate differences in memory retrieval and language processing between high-IQ and low-IQ groups. Similarly, occipital regions, particularly [O2], revealed distinctions in visual-spatial processing capabilities. At the same time, non-significant channels like [P3] and [PZ] in the posterior parietal regions, [T6] in the temporal region and [F7] in the left frontal area pointed to areas where further exploration or alternative modeling approaches may be necessary.

The low-IQ group exhibited higher predictability during Monte Carlo simulations, as demonstrated by lower RMSE values between real and simulated data and smaller standard deviations in several EEG channels. The less intelligent subjects generally displayed more consistent EEG patterns, particularly in central and temporal regions, aligning with their reduced variability and narrower distribution ranges. These patterns suggest a level of neural uniformity in the low-IQ group, contrasting with the broader variability, the higher RMSE values and the increased unpredictability observed in high-IQ individuals’ EEG activity.

We demonstrated that Monte Carlo simulations serve as a powerful tool in neurocognitive research, enabling the identification of EEG markers associated with cognitive abilities and the simulation of neural activity patterns to test hypotheses about brain function. Practical applications of these findings extend to predictive modeling, where simulated EEG data can forecast cognitive behaviors. In clinical settings, these insights might inform interventions for cognitive enhancement or rehabilitation. Additionally, educational strategies could be tailored based on neural markers of learning potential. This study has limitations. While the assumption of normality proved valid for most EEG channels, deviations in skewness or kurtosis in some columns may have influenced results, highlighting the need for further research. Future studies should consider employing non-parametric methods or fitting alternative distributions to enhance the robustness of simulations. Investigating multivariate correlations across channels could provide deeper insights into the neural interconnections underlying intelligence. Future research should also explore the integration of alternative modeling techniques to further refine the accuracy and applicability of these methods and deepen our understanding of the neural underpinnings of human cognition.

In conclusion, the implications of this research in the field of cognitive neuroscience go beyond academic interest, underscoring the potential of combining advanced statistical techniques with neuroscience to unlock new possibilities for studying and enhancing human cognition. In particular, Monte Carlo simulations leverage the power of computational modeling to explore the complexities of human intelligence, offering robust methods for expanding datasets, uncovering underlying patterns and identifying key neural markers.

## DECLARATIONS

### Ethics approval and consent to participate

This research does not contain any studies with human participants or animals performed by the Author.

### Consent for publication

The Author transfers all copyright ownership, in the event the work is published. The undersigned author warrants that the article is original, does not infringe on any copyright or other proprietary right of any third part, is not under consideration by another journal, and has not been previously published.

### Availability of data and materials

all data and materials generated or analyzed during this study are included in the manuscript. The Author had full access to all the data in the study and take responsibility for the integrity of the data and the accuracy of the data analysis.

## Acknowledgements

none.

## Authors’ contributions

The Author performed: study concept and design, acquisition of data, analysis and interpretation of data, drafting of the manuscript, critical revision of the manuscript for important intellectual content, statistical analysis, obtained funding, administrative, technical, and material support, study supervision.

## Declaration of generative AI and AI-assisted technologies in the writing process

During the preparation of this work, the author used ChatGPT to assist with data analysis and manuscript drafting. After using this tool, the author reviewed and edited the content as needed and takes full responsibility for the content of the publication.

